# Reconstruction Set Test (RESET): a computationally efficient method for single sample gene set testing based on randomized reduced rank reconstruction error

**DOI:** 10.1101/2023.04.03.535366

**Authors:** H. Robert Frost

## Abstract

We have developed a new, and analytically novel, single sample gene set testing method called Reconstruction Set Test (RESET). RESET quantifies gene set importance at both the sample-level and for the entire dataset based on the ability of set genes to reconstruct values for all measured genes. RESET addresses four important limitations of current techniques: 1) existing single sample methods are designed to detect mean differences and struggle to identify differential correlation patterns, 2) computationally efficient techniques are self-contained methods and cannot directly detect competitive scenarios where set genes differ from non-set genes in the same sample, 3) the scores generated by current methods can only be accurately compared across samples for a single set and not between sets, and 4) the computational performance of even the fastest existing methods be significant on very large datasets. RESET is realized using a computationally efficient randomized reduced rank reconstruction algorithm (available via the RESET R package on CRAN) that can effectively detect patterns of differential abundance and differential correlation for self-contained and competitive scenarios. As demonstrated using real and simulated scRNA-seq data, RESET provides superior accuracy at a lower computational cost relative to other single sample approaches.

## 1 Introduction

### 1.1 Gene set testing

High-dimensional genomic profiling technologies, such as RNA-sequencing, give researchers a powerful, molecular-level picture of tissue and cellular biology, however, the gain in fidelity obtained by measuring thousands of genomic variables comes at the price of impaired interpretation, loss of power due to multiple hypothesis correction and poor reproducibility [1, 2]. To address these challenges for bulk tissue data, researchers developed gene set testing, or pathway analysis, methods [3–6]. Gene set testing is a widely used and effective hypothesis aggregation technique that analyzes biologically meaningful groups of genes, e.g., the genes involved in specific signaling pathway defined in a resource like Reactome [7], instead of individual genomic variables. Focusing on a collection of pathways can significantly improve power, interpretation and replication relative to an analysis focused on individual genes [3, 8]. The benefits of gene set testing are even more pronounced for single cell data given increased technical variance and sparsity [9, 10].

Gene set testing methods can be grouped according to four main features:

1. *Supervised vs unsupervised*: Does the method test for the association between gene set members and a specific outcome or does it generate gene set scores using only the measured genomic data?
2. *Population vs single sample*: Does the method generate gene set scores for each sample or just a single score for the entire population?
3. *Self-contained vs competitive*: Does the method test the *H*_0_ that none of the genes in the set has an association with the outcome or the *H*_0_ that the genes in the set are not more associated with the outcome than genes not in the set?
4. *Uniset vs multiset*: Does the method test each gene set separately (uniset) or jointly evaluate all sets in a collection (multiset)?

The most popular type of gene set test is uniset, population-based, competitive and supervised, which is driven by several factors: 1) uniset tests are easier to implement and execute than multiset tests, 2) biological hypotheses of interest typically correspond to supervised tests (e.g., differential expression relative to a specific clinical variable), and 3) a competitive *H*_0_ often generates more meaningful results than a self-contained *H*_0_ [8]. Although gene set analysis can be performed on a variety of omics data types, it is most commonly applied to transcriptomics data and, without loss of generality, we will assume this data type in the remainder of the manuscript.

### 1.2 Single sample gene set testing

Although supervised and population-level methods such as GSEA [2] and CAMERA [11] are the most commonly used gene set testing techniques, unsupervised and single sample methods have become increasingly popular given their significant analytical flexibility. Single sample methods, which are inherently unsupervised, operate like a variable transformation to convert an input *n × p* matrix **X** that captures expression of *p* genes in *n* samples into an *n × m* matrix **S** that captures the sample-level enrichment of *m* gene sets. This matrix of sample-level gene set scores can then be used in a wide range of subsequent computational tasks including unsupervised analyses like data visualization and supervised analyses like testing the association of each column of **S** with a given outcome variable, which generates results similar to those created by a population-level and supervised technique.

A number of single sample gene set testing methods are currently available, which can be generally grouped into self-contained and competitive categories. Competitive techniques like GSVA [12] and ssGSEA [13]) generate sample-level scores using a Kolmogorov-Smirnov (KS) like random walk statistic computed on the gene ranks within each sample, often following some form of gene standardization across the samples. Self-contained methods like PLAGE [14], PAGODA [15], the technique of Lee et al. [16], scSVA [17], Vision [18], and our VAM method [9] generate scores using only the data for genes in the set. Our development of the VAM technique was motivated by the poor performance of other single sample techniques on single cell transcriptomic data. Specifically, we found that existing techniques have poor classification performance in the presence of sparsity and technical noise, and a high computational cost. The VAM method is a novel modification of the standard Mahalanobis multivariate distance measure that generates cell-specific gene set scores which account for the inflated noise and sparsity of single cell RNA-sequencing (scRNA-seq) data. Because the distribution of the VAM-generated scores has an accurate gamma approximation under the null of uncorrelated technical noise, these scores can also be used for inference regarding pathway activity.

### 1.3 Single sample gene set testing challenges

While the VAM technique offers a significant improvement in terms of computational performance and accuracy over other single sample methods, it has four important limitations:

1. All existing single sample gene set testing methods are designed to detect differences in mean value and struggle to identify biologically relevant patterns of differential correlation.
2. The VAM method, and other computationally efficient techniques like PLAGE [14] and the z-scoring method of Lee et al. [16], are self-contained methods that generate scores for a given gene set without considering the values of genes not in the set. These self-contained methods cannot directly detect competitive scenarios where the measured values of set genes differ from non-set genes in the same sample.
3. The scores generated by existing single sample methods can only be accurately compared across samples for a single set and not between sets. This limitation complicates many types of multivariate downstream analyses that attempt to jointly evaluate the scores for multiple sets. For example, it becomes challenging to determine which of several gene sets are more active/enriched in a given sample since the scores for different sets are not necessarily on the same scale.
4. The computational performance of VAM, while better than most existing methods, can still be significant on very large datasets. For VAM, and other computationally efficient self-contained methods, computational cost directly scales with the number of samples. As the price of single cell experimental methods continues to fall, the number of cells in a typical dataset has grown substantially with tens-to-hundreds-of-thousands of cells now common. Projects that generate single cell data on samples from hundreds of separate patients will result in even larger total sample sizes. These very large single cell datasets motivate performance improvements beyond what can be obtained using existing techniques like VAM.

### 1.4 Gene set testing based on reconstruction error

To address the challenges faced by VAM and other single sample approaches, we have developed a new, and analytically novel, single sample gene set testing method called Reconstruction Set Test (RESET). RESET quantifies gene set importance at the sample-level and for the entire input data matrix based on the ability of genes in the set to reconstruct values for all measured genes. RESET is realized using a computationally efficient randomized reduced rank reconstruction algorithm and can effectively detect patterns of differential abundance and differential correlation for both self-contained and competitive scenarios. The use of reconstruction error by RESET is quite distinct from standard approaches to gene set testing and has the potential to capture biological patterns not detectable using methods based on differences in mean expression. Unique among single sample methods, RESET generates both overall and sample-level scores for evaluated gene sets. Mathematical details of the RESET method and the evaluation design are outlined in Section 2. Section 3 contains the simulation study and real data analysis results, which demonstrate that RESET provides superior classification accuracy at a lower computation cost relative to VAM and other popular single sample gene set testing approaches. An R implementation, which supports integration with the Seurat framework, is available in the RESET package on CRAN.

## 2 Methods

### 2.1 Reconstruction Set Test (RESET) overview

The RESET method computes sample-specific and overall gene set scores from gene expression data using the error from a randomized reduced rank reconstruction. At a high-level, RESET takes as input two matrices:

- **X**: *n × p* matrix that holds the abundance measurements for *p* genes in *n* samples.
- **A**: *m × p* matrix that represents the annotation of the *p* genes in **X** to *m* gene sets as defined by a collection from a repository such as the Molecular Signatures Database (MSigDB) [19] (*a*_*i,j*_ = 1 if gene *j* belongs to gene set *i*).

RESET generates as output:

- **S**: *n × m* matrix that holds sample-specific gene set scores for each of the *n* samples in **X** and *m* gene sets defined in **A**.
- v: length *m* vector that holds the overall scores for each of the *m* gene sets defined in **A**.

The version of the RESET method implemented in the RESET R package and used to generate the results in Section 3 accepts a number of additional parameters (center, scale, num.pcs, pca.buff, pca.q, random.threhold, k, k.buff, q, test.dist, norm.type) whose function, motivation and interdependencies are fairly complex. To make the full method easier to understand, we will start by defining a simplistic, and computationally inefficient, version of RESET in Section 2.2, refine that version to use a more efficient reduced rank reconstruction in Section 2.3, before detailing the fully optimized implementation that incorporates randomized numerical linear algebra (RNLA) methods in Section 2.4.

### 2.2 Simplistic RESET

A simplistic version of the RESET method is detailed in Algorithm 1 below. This version of RESET uses all of the genes in each set to reconstruct the full matrix **X**, generates overall scores using the Frobenius norm of the reconstruction error matrix, and generates sample-level scores using the Eucledian norm of the reconstruction error for the associated row.

#### Algorithm 1

Simplistic RESET

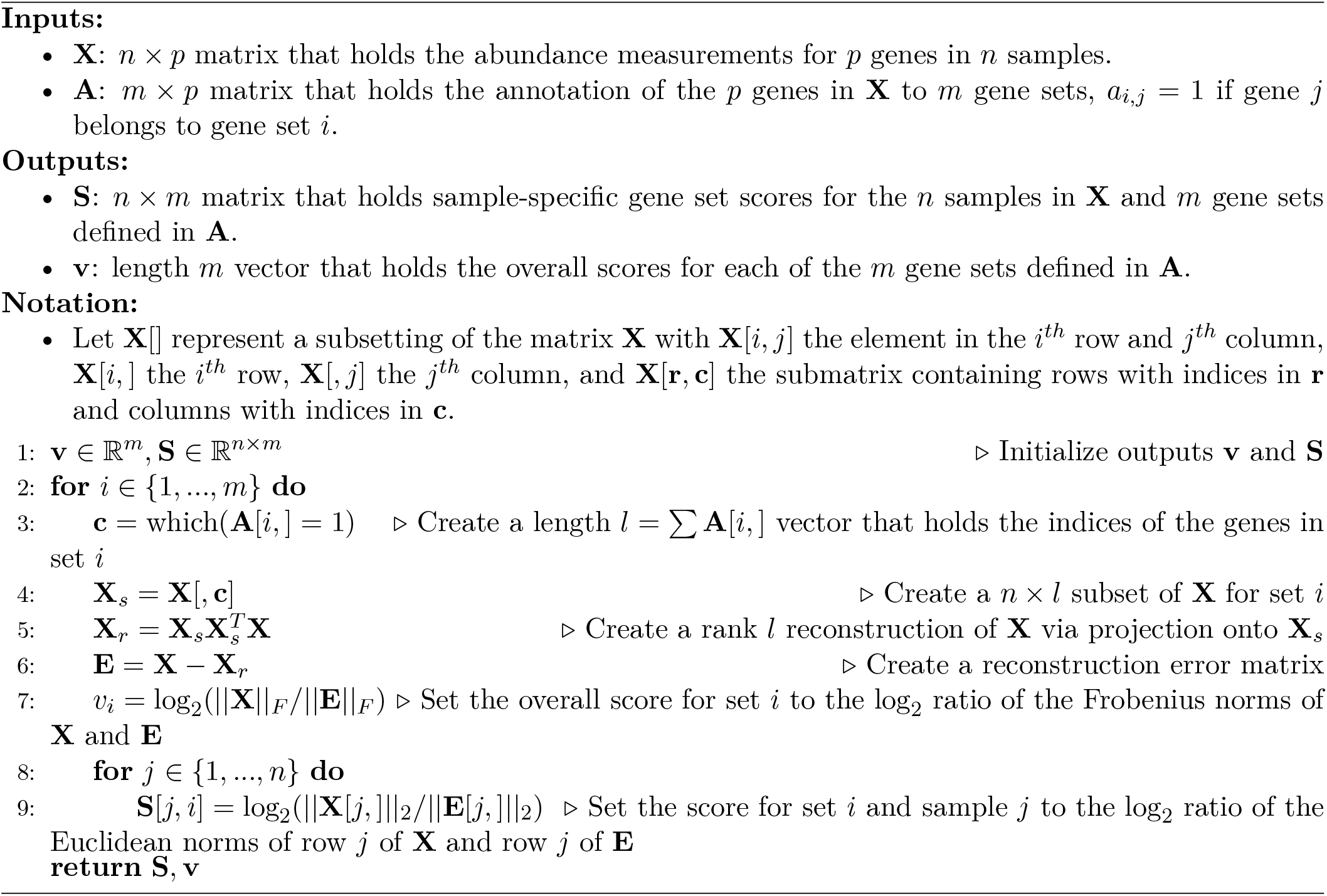

Although the simplistic version of RESET captures the general structure of the method, it has several critical limitations:

- Computational cost can be significant if either **X** or the gene sets defined in **A** are large.
- Reconstruction of the full **X** matrix using all genes in a given set can produce scores in **v** and **S** that are dominated by noise when the biological signal in **X** has an effective rank that is much lower than the observed rank of **X**, which is common for genomic data.
- If gene sets defined by **A** have distinct sizes, the generated scores under a null scenario of completely random data in **X** will not be equivalent. In particular, scores will be elevated for larger sets as compared to smaller sets.

### 2.3 Reduced rank RESET

The limitations of the simplistic version of RESET defined in Algorithm 1 can be effectively addressed by reconstructing a dimensionally reduced version of **X** using a dimensionally reduced version of **X**_*s*_ (the subset of **X** corresponding to each gene set). This approach can be efficiently realized by projecting **X** onto the top *b* principal components (PCs) of **X** where *b* is close to the rank of the biological component of the data and then assessing how well this PC projection can be reconstructed using a rank *k* basis for the column space of each **X**_*s*_. This reduced rank version of RESET is defined in Algorithm 2 below.

#### Algorithm 2

Reduced rank RESET

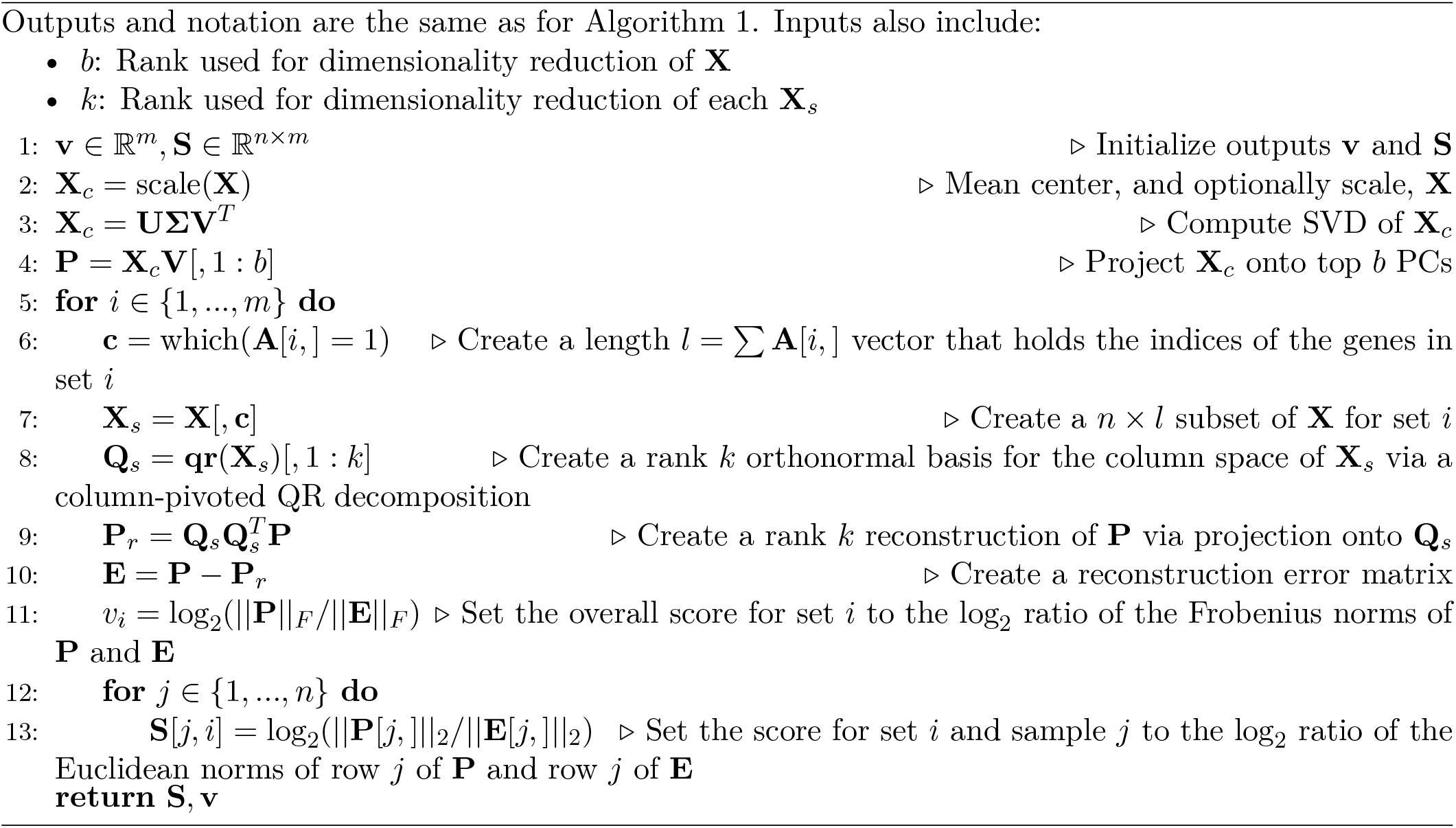

Outputs and notation are the same as for Algorithm 1. Inputs also include:

### 2.4 Randomized RESET

Although the reduced rank version of RESET detailed in Algorithm 2, successfully addresses the key limitations of the simplistic version, computational cost can still be significant. In particular, the PCA and QR computation steps can be very expensive, even if truncated algorithms are used that halt after computing the top PCs/columns (e.g., the truncated PCA algorithm implemented in the irlba R package [20] or a truncated column-pivoted QR decomposition). Fortunately, the computational performance of these matrix decompositions can be dramatically improved by leveraging randomized numerical linear algebra (RNLA) [21, 22] techniques with only minimal loss of accuracy. Such RNLA methods have been successfully leveraged for the analysis of large genomic data matrices, e.g., scRNA-seq data, with data imputation via reduced rank reconstruction a key use case [23,24]. Before detailing the randomized RESET algorithm, we need to define two underlying RNLA functions: a randomized technique for computing an orthonormal basis for the column space of a matrix and, building on that method, a randomized SVD algorithm. These two RNLA techniques follow the general structure of the randomized rangefinder and randomized SVD algorithms in Martinsson et al. [21]. For clarity, we are not including power iteration functionality in Algorithms 3 or 4 (see Martinsson et al [21] or Erichson et al. [25] for details on power iteration functionality). We are also showing just the use of *N* (0, 1) random variables for creation of the sketch matrix. While the RESET method supports both *N* (0, 1) and *U* (0, 1) RVs, empirical studies have found that performance of randomized methods is generally insensitive to the choice of statistical distribution [21]. Readers interested in the theoretical and computation properties of these methods or the broader foundations/applications of RNLA are encouraged to read the excellent survey by Martinsson et al. The paper by Erichson et al [25] associated with the *rsvd* R package provides a shorter introduction to these methods with a specific focus on their programmatic implementation and performance benefits relative to truncated algorithms.

#### Algorithm 3

Randomized column space basis generator (randomColumnSpace)

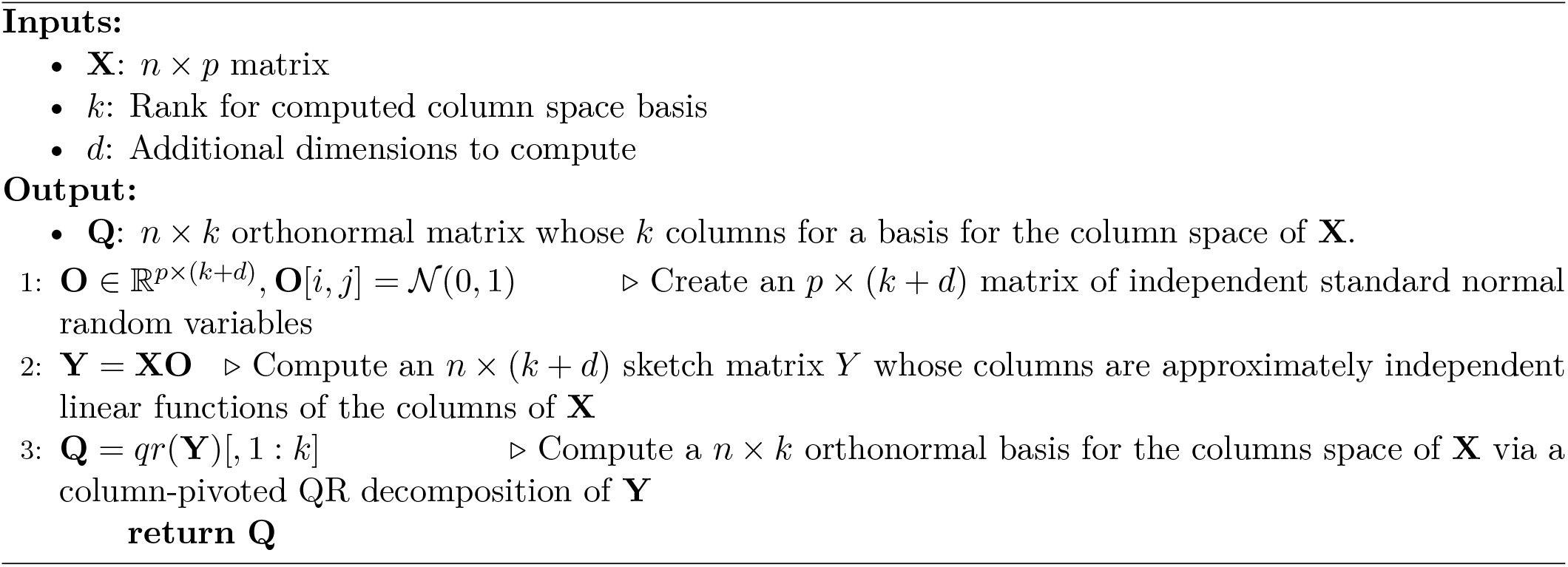

#### Algorithm 4

Randomized SVD (randomSVD)

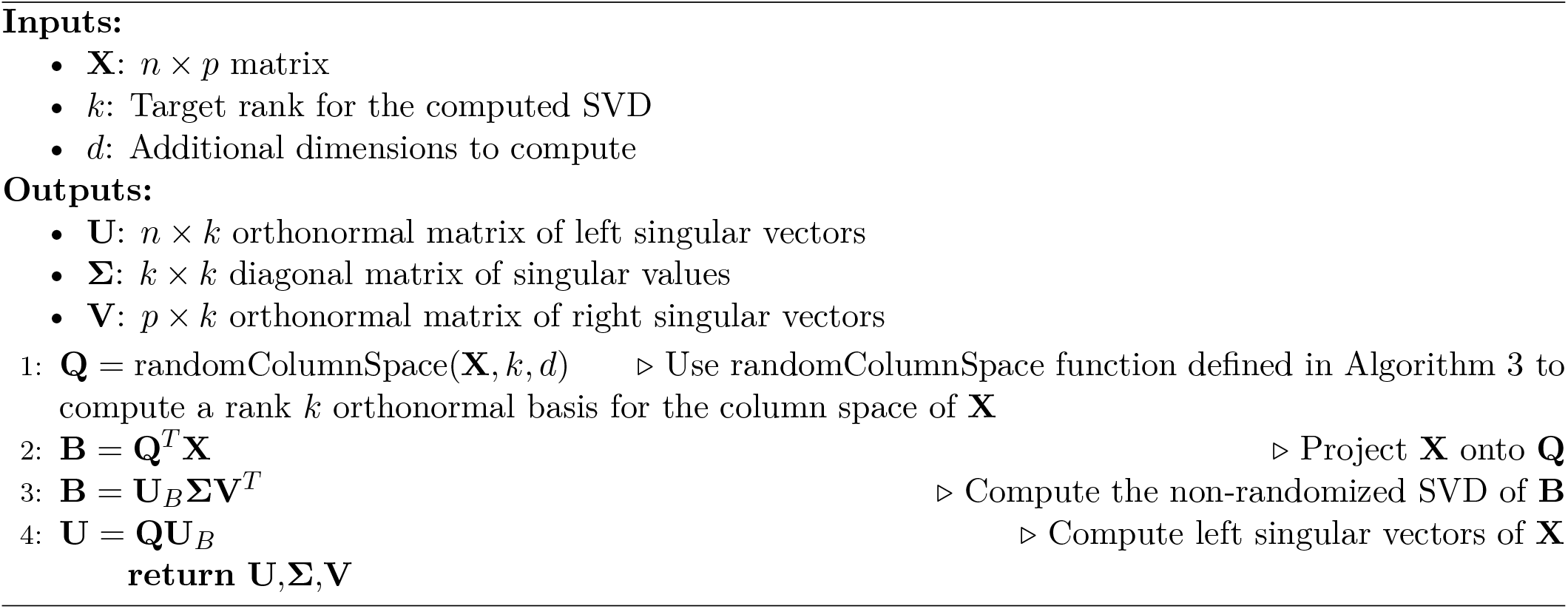

Given the randomized column space basis generator defined in Algorithm 3 and the randomized SVD defined in Algorithm 4, we can now detail the randomized version of RESET. For clarity, we are omitting parameters that allow for delayed mean centering of **X** (see Section 2.6 and the RESET R package documentation for more details).

#### Algorithm 5

Randomized RESET

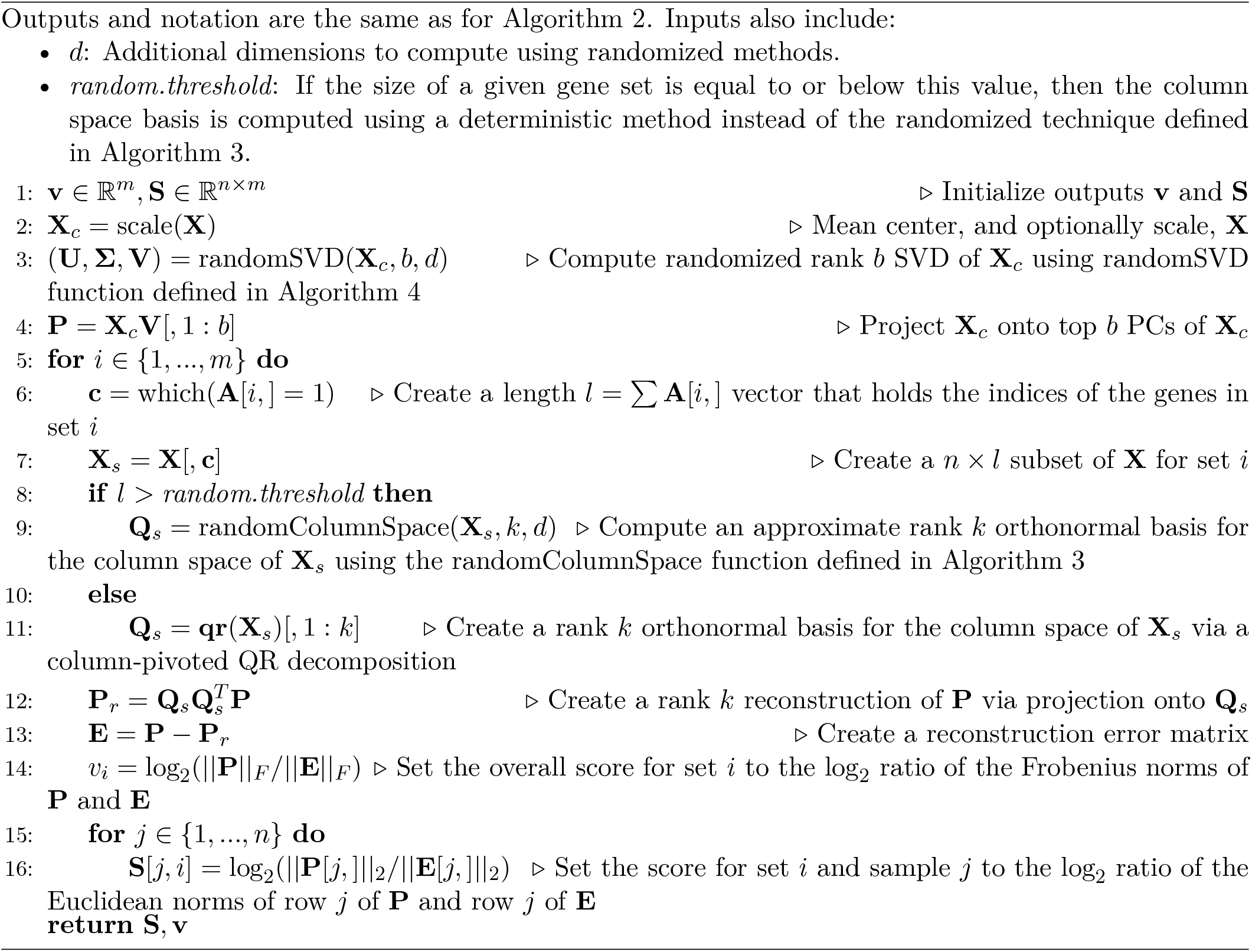

Outputs and notation are the same as for Algorithm 2. Inputs also include:

### 2.5 RESET-Seurat integration

To support the analysis scRNA-seq data, the RESET R package includes a wrapper function (*resetForSeurat()*) that enables direct integration with the popular Seurat framework [26]. This integration supports both log-normalization and SCTransform [27] normalization and assumes that PCA has already be performed via the Seurat *RunPCA()* function. To use an alternative data subset or transformation for measuring reconstruction error, the underlying *reset()* function must be called directly. The **S** matrix of cell-level gene set scores output by RESET is saved as a new Seurat assay named “RESET”, which enables the visualization and further analysis of these scores using Seurat framework, e.g., the *FeaturePlot()* and *FindMarkers()* functions. The vector **v** of overall gene set scores is saved in the feature metadata column named “RESET”. See the RESET R package documentation and vignettes for further details and examples.

### 2.6 RESET usage considerations

The randomized RESET method defined in Algorithm 5 and implemented in the RESET R package supports a number of parameters that enable customization of the method for different analysis scenarios. Two important use cases are the application of RESET to large sparse data sets and evaluation of gene set collections that contain sets whose size is close to the target rank *k*. Considerations for these scenarios, the selection of appropriate values for *k* and *b*, and deciding how to leverage the generated **S** and **v** scores, are discussed below.

- **Sparse X:** When **X** is large and sparse, which is typical of single cell data, it is typically represented using an optimized sparse matrix format (e.g., the sparse matrix support in the R *Matrix* package). In this case, mean centering of the columns of **X** will force conversion into a dense matrix format, which can have a significant impact on both memory usage and computational complexity for subsequent matrix operations. To avoid this performance penalty, it desirable to only mean center a subset of **X** containing the data needed for gene set testing. How this scenario can be handled for RESET depends on whether the method is being executed via the Seurat framework interface (i.e., the *resetForSeurat()* function) or directly via either the *resetViaPCA()* or *reset()* functions. In both cases, RESET can be executed such that mean centering is only applied to a subset of **X** containing the genes that belong to the evaluated gene sets, i.e., it does not force the entire **X** matrix into a dense format. When RESET is executed via the Seurat framework interface, it is assumed that PCA has already been performed on a scaled and mean centered version of the normalized scRNA-seq data. Because Seurat by default only applies mean centering to a subset of the scRNA-seq data corresponding to the genes with the largest biological variance, the memory and performance impact is less severe. The Seurat wrapper passes in the unscaled normalized scRNA-seq matrix to the *reset()* function with parameters set so that mean centering of **X** is only performed on the columns for each gene set. If RESET is executed via the *resetViaPCA()* function, this delayed mean centering can be enabled by setting the *center* parameter to false, which will result in **X** being projected onto the uncentered PCs and centering performed on just the PC projections and subsets of **X** corresponding to each gene set. If RESET is executed via the *reset()* function, then users have full control over mean centering behavior (see the R package documentation and vignettes for more details).
- **Gene set size is close to target** *k*: The computational benefit of the randomized column space basis generator detailed in Algorithm 3 is only meaningful if the number of columns in the input matrix is at least 3 times larger than the target rank *k* [25]. This means that use of randomization for gene sets whose size is less than *∼* 3*k* will incur an accuracy penalty without any improvement in execution time. In this case, it is desirable to instead use the column-pivoted QR decomposition approach. The randomized RESET method detailed in Algorithm 5 supports this flexibility via the *random*.*threshold* argument. Randomization can be required for all evaluated sets by setting *random*.*threshold* to a value that is less than the minimal gene set size. Similarly, use of the deterministic column-pivoted QR decomposition for all gene sets can be achieved by setting *random*.*threshold* to the maximum gene set size.
- **Selecting appropriate values for** *b* **and** *k*: How to select the target rank for reduced rank matrix decompositions is a long standing problem in applied mathematics. Although the RESET method does not directly address the rank selection problem and leaves specification of these parameters to users, there are a number of established approaches that can be followed when determining appropriate values for *b* and *k*. Most of these techniques select the target rank based on the distribution of singular values using either a heuristic criteria, e.g., the elbow method, a model-based threshold, e.g., use of the random matrix theory-based eigenvalue null distribution [28], or a resampling technique, e.g., the JackStraw procedure used in the Seurat framework [29]. The simulation and real data results presented in Section 3 use approximate, and likely non-optimal, values for *b* and *k*; performance of RESET in these cases could probably be improved through use of a more sophisticated rank selection method.
- **Using the S and v scores:** Unique among single sample gene set testing methods, RESET generates both sample-level scores in the **S** matrix and overall scores in the **v** vector. The sample-level scores provide the most flexibility and can be used in place of gene abundance data in a wide range of subsequent statistical analysis, e.g., differential expression analyses, regression modeling, clustering, visualization, etc. Although the overall gene set scores offer less general utility, they still provide distinct information regarding the overall biological signal in the data and can be leveraged for filtering or weighting of pathway-based models.

### 2.7 Simulation study design

To evaluate the performance of the RESET method, we applied it to simulated sparse count data with statistical charateristics similar to those found in scRNA-seq data. In particular, we simulated **X** matrices representing normalized scRNA-seq data for 2,000 cells and 500 genes as follows:

- Populate all entries of **X** with independent negative binomial random variables with a mean of 0.2 and overdispersion of 100. The mean value of 0.2 is based on the mean value for unnormalized counts in the PBMC3k scRNA-seq data used for the real data analysis (see Section 2.8 for details) and the overdispersion value of 100 is based on the finding by Lause et al. [30] that scRNA-seq counts can be effectively modeled by a negative bionomial distribution with flexible mean and fixed overdispersion of 100.
- Modify the counts for a subset of the entries in **X** corresponding to a hypothetical gene set to be correlated with an inflated mean. We refer to this scenario, which generates a pattern that can be detected by both self-contained and competitive methods, as the “block” design. The number of genes in the hypothetical set was varied between 10 and 90 and the number of cells with modified counts for the set was varied between 25 and 425. The mean for these entries was varied between 0.2 (the no differential expression case) and 0.44 and the inter-gene correlation was varied between 0 (the no differential correlation case) and 0.9. Two special scenarios were also simulated: the pure self-contained case and the pure competitive case. For the pure self-contained case, all 500 genes were included in the set. For the pure competitive case, counts were modified for all 2,000 cells. A total of 50 **X** matrices were simulated for each unique combination of simulation parameters for each of the scenarios.
- Perform log-normalization of the count data to mirror Seurat’s log-normalization procedure, i.e., divide the counts by the total for each cell, multiply by a scale factor of 10,000, add a pseudocount of 1 and take the natural log.

These simulated **X** matrices were used to evaluate the classification and computational performance RESET and the comparison methods detailed in Section 2.9. Two different versions of RESET, RESET.det and RESET.ran, were evaluated based on the setting of the *random*.*threshold* parameter. For RESET.det, which stands for deterministic RESET, *random*.*threshold* was set to 90, which forces the use of column-pivoted QR decomposition. For RESET.ran, which stands for randomized RESET, *random*.*threshold* was set to 9, which forces the use of the randomized column space basis generator. Both RESET.ran and RESET.det were realized by calling the *resetViaPCA()* function in version 0.1.0 of the RESET R package with the following settings for all parameters except *random*.*threshold*: *num*.*pcs=10, pca*.*buff=10, pca*.*q=2, k=10, k*.*buff=0, q=0, test*.*dist=”normal”, norm*.*type=“2”*. Single sample gene set scores were computed using each evaluated technique for all simulated **X** matrices and AUC values were calculated based on the ability of each method to give modified cells a higher score than unmodified cells. Overall classification performance was also assessed for the two RESET versions (other methods could not be evaluted since they do not generate overall scores). To assess overall classification performance, a total of five equally sized sets were scored with only the first set containing modified counts. The same five set design was also used to assess classification performance for the pure competitive scenario, i.e., single sample scores for all five sets were included in the AUC computation. Note that execution time for RESET did not include generation of the PC projections via randomized SVD since that is only called once for all evaluated gene sets so would disporportionately impact the results when only a single gene set is evaluated. The time required for computing the PC projections is also not a factor for Seurat-based analyses since PCA is typically performed regardless for clustering and visualization.

### 2.8 Real data analysis design

To evaluate RESET on real transcriptomic data, we analyzed two public scRNA-seq data sets available from 10x Genomics: the 2.7k human PBMC data set used in the Seurat Guided Clustering Tutorial [31], and an 11.8k mouse brain cell data set generated on the combined cortex, hippocampus and sub ventricular zone of an E18 mouse [32]. These data sets were selected in part because they were also used in the VAM paper so provide a direct measure of comparative performance. Similar to the rationale listed in the VAM paper, these data sets are representative of small and medium sized experiments and capture gene expression profiles for two different complex cells populations (immune cells and neural cells) from two organisms (human and mouse) that comprise a large percentage of existing scRNA-seq data. Preprocessing, quality control (QC), normalization and clustering of the PBMC data set matched the Seurat Guided Clustering Tutorial. Specifically, the Seurat log-normalization method is used followed by application of the *vst* method for decomposing technical and biological variance. Preprocessing and QC of the PBMC data yielded an **X** matrix of normalized counts for 14,497 genes and 2,638 cells.

Processing of the mouse brain data followed similar quality control metrics (at least 200 features per cell, non-zero values in at least 10 cells for genes, proportion of mitochondrial reads less than 10% [33]) with Uniform Manifold Approximation and Projection (UMAP) [34] used for dimensionality reduction and clustering performed with Seurat’s implementation of shared nearest neighbor (SNN) modularity optimization [35]. Normalization of the mouse brain data was performed using SCTransform [27] rather than log-normalization to assess RESET performance for both of the supported Seurat normalization approaches. Preprocessing and QC of the mouse brain data yielded an **X** matrix of normalized counts for 32,850 genes and 9,320 cells.

For these analyses, the gene set matrix **A** was populated using the human C2.CP.BIOCARTA (BioCarta, 292 gene sets), and the mouse C5.BP (Gene Ontology Biological Processes, 7,751 gene sets) collections from v2023.1 of the Molecular Signatures Database (MSigDB) [19]. These MSigDB collections represent two widely used groups of curated gene sets: BioCarta [19], and the biological process branch of the Gene Ontology [36]. Prior to performing gene set testing, the Entrez gene IDs used by MSigDB were converted to Ensembl IDs using logic in the Bioconductor *org*.*Hs*.*eg*.*db* and *org*.*Mm*.*eg*.*db* R packages. The **X** and **A** matrices were then filtered to only contain genes present in both matrices (13,714 genes for the PBMC data and 16,425 genes for the mouse brain data). Finally, the **A** matrix was filtered to remove sets with fewer than 5 or more than 200 members. Enrichment of gene sets for specific scRNA-seq clusters was performed using a Wilcoxon rank sum test as implemented by the Seurat *FindMarkers()* method.

Execution of RESET for both of these data sets was performed using the *resetForSeurat* method in v0.1.0 of the RESET R package with the following parameter settings: *num*.*pcs=15, k=5, k*.*buff=0, q=0, random*.*threshold=30, test*.*dist=”normal”, norm*.*type=“2”*.

### 2.9 Comparison methods

To assess the relative performance of the RESET method on simulated and real scRNA-seq data, we used our previous developed VAM method [9] and two other popular single sample techniques: GSVA [12] and ssGSEA [13]. For VAM, we used the implementation in version 1.0.0 of the VAM R package from CRAN. For GSVA and ssGSEA, we used the implementations available in version 1.46.0 of the GSVA R package from Bioconductor. Unless otherwise noted, all the comparison methods were executed using default parameter values.

## 3 Results and discussion

### 3.1 Sample-level classification performance

To compare the performance of RESET against our previously developed VAM technique and the popular GSVA and ssGSEA methods, we measured the classification accuracy (i.e., the ability of each method to generate high scores for cells that have higher mean expression and/or non-zero correlation between genes in a specific set) on scRNA-seq data sets simulated following the designs detailed in Section 2.7. Figure 1 illustrates the relative classification performance (as measured by the area under the receiver operating characteristic curve (AUC)) of RESET.det, RESET.ran, VAM, GSVA [12], and ssGSEA [13] for the block design across a range of mean inflation, inter-gene correlation, gene set size and number of informative samples (i.e., number of cells for which gene set values are inflated and/or correlated). Figures 2 and 3 provide similar results for the pure self-contained and pure competitive scenarios. All of these figures display the average AUC (and standard error of the mean via error bars) for 50 simulated data sets generated for each distinct combination of parameter values. Importantly, both the deterministic and randomized versions of the RESET method provide superior classification accuracy relative to VAM, GSVA and ssGSEA across nearly the full range of evaluated parameter values for all three simulation designs with significant relative performance benefits for the pure self-contained and pure competitive cases.

**Figure 1:**
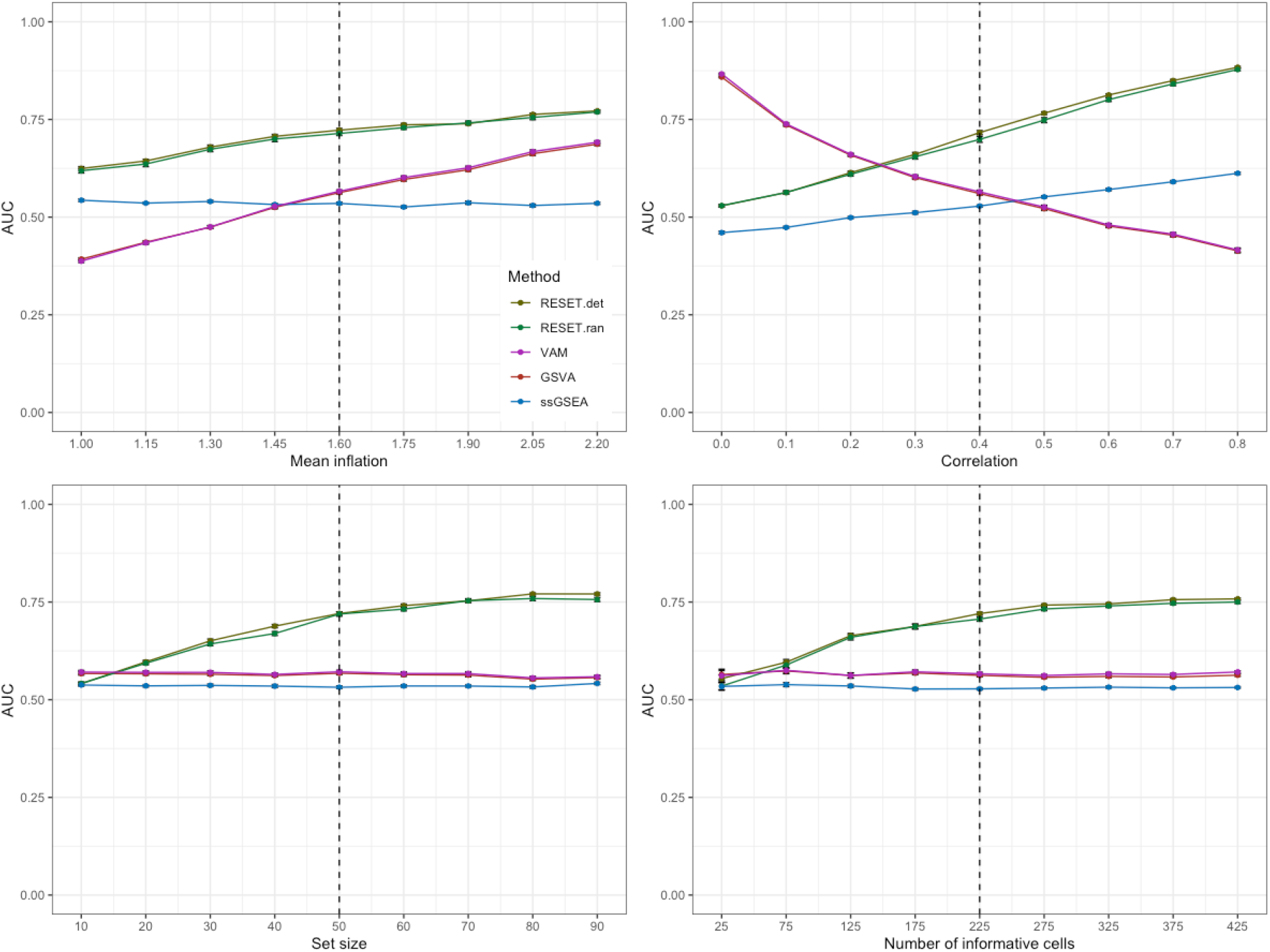
Classification performance of RESET.det, RESET.ran, VAM, GSVA, and ssGSEA on scRNA-seq data simulated according to Section 2.7 for the block design. Each panel illustrates the relationship between the area under the receiver operating characteristic curve (AUC) and one of the simulation parameters. The vertical dotted lines mark the default parameter value used in the other panels. Error bars represent the standard error of the mean.

**Figure 2:**
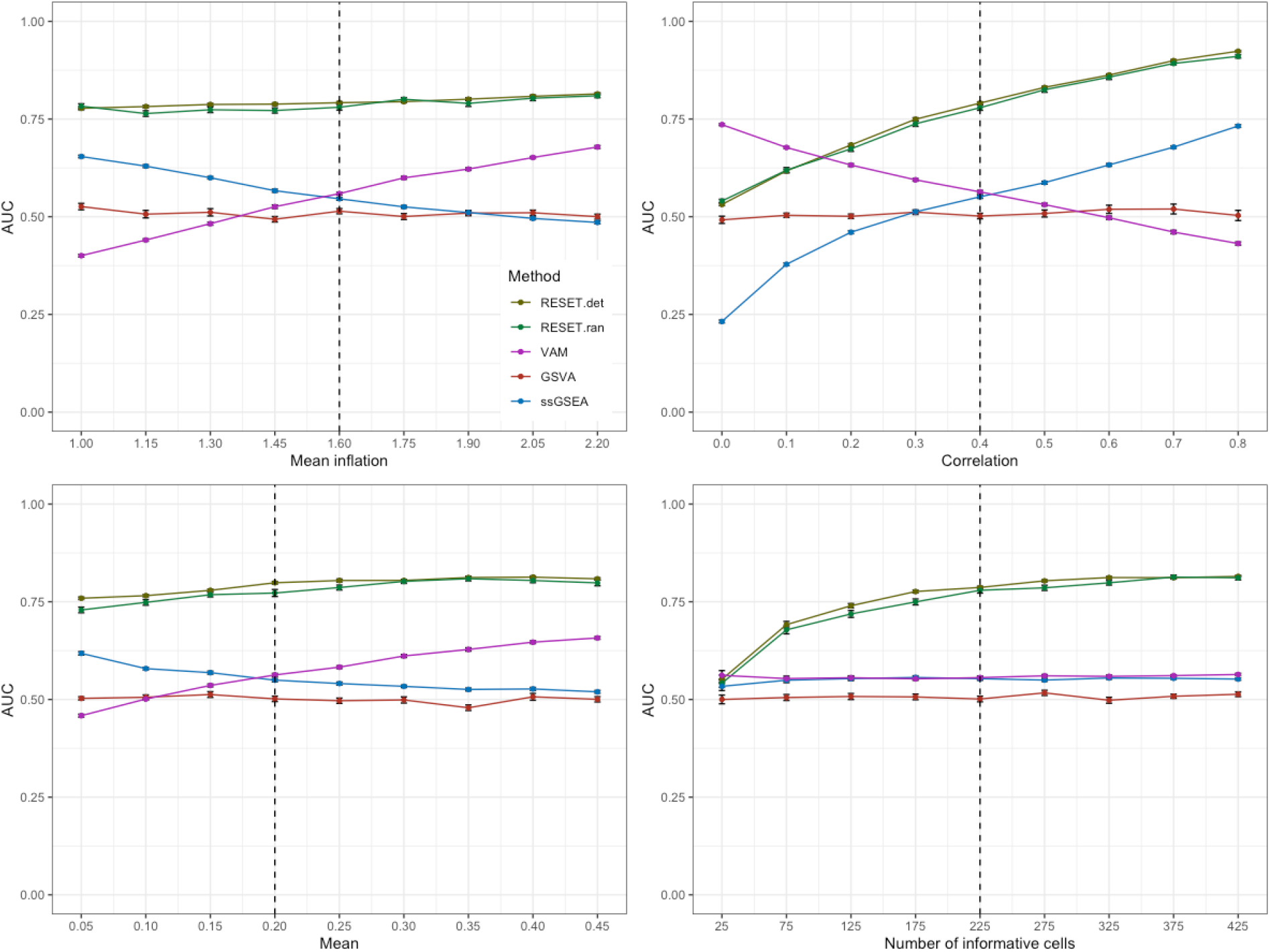
Classification performance of RESET.det, RESET.ran, VAM, GSVA, and ssGSEA on scRNA-seq data simulated according to Section 2.7 for the pure self-contained design. Each panel illustrates the relationship between the area under the receiver operating characteristic curve (AUC) and one of the simulation parameters. The vertical dotted lines mark the default parameter value used in the other panels. Error bars represent the standard error of the mean.

**Figure 3:**
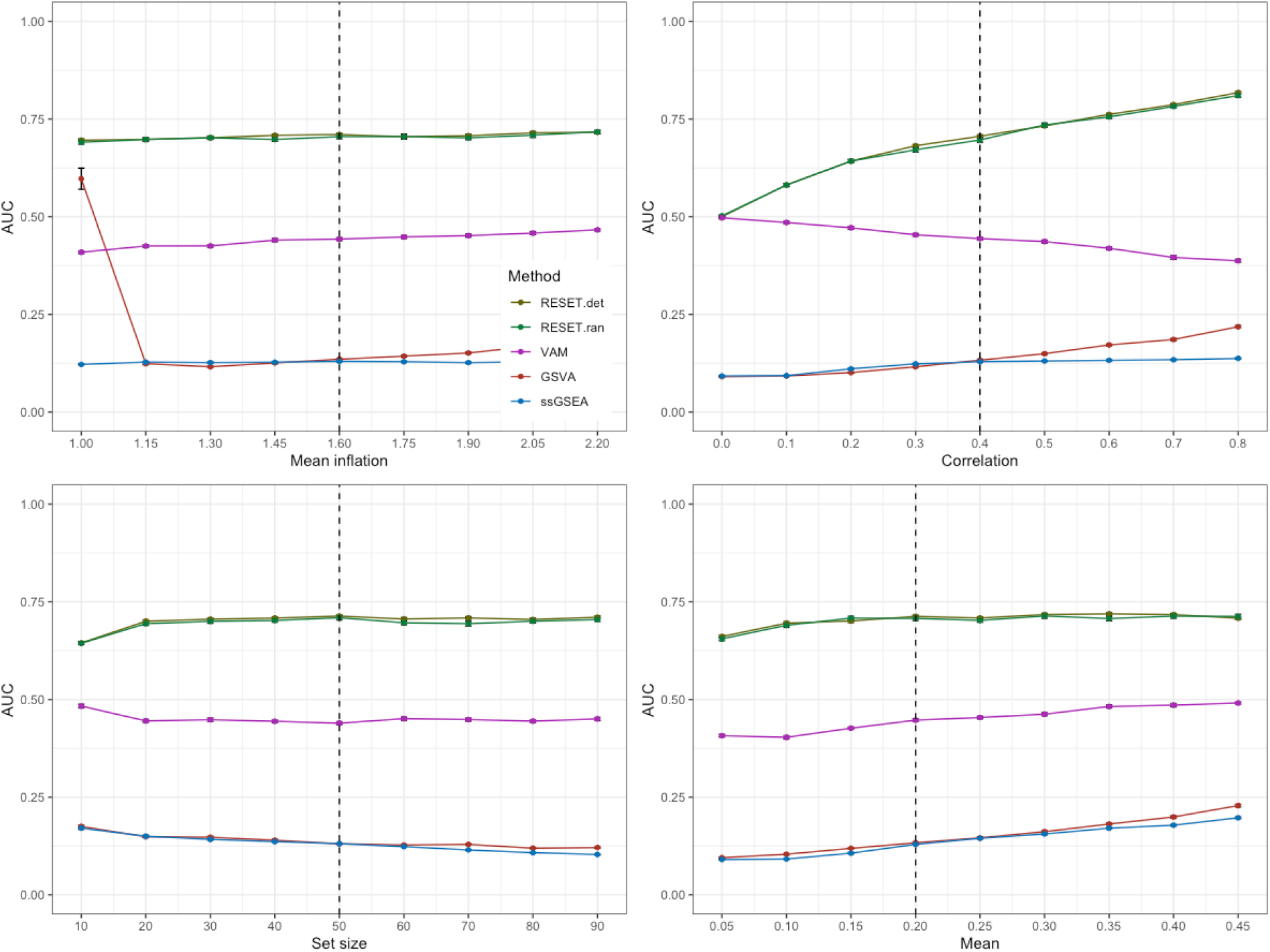
Classification performance of RESET.det, RESET.ran, VAM, GSVA, and ssGSEA on scRNA-seq data simulated according to Section 2.7 for the pure competitive design. Each panel illustrates the relationship between the area under the receiver operating characteristic curve (AUC) and one of the simulation parameters. The vertical dotted lines mark the default parameter value used in the other panels. Error bars represent the standard error of the mean.

The results for the block simulation design visualized in Figure 1 indicate that the two versions of RESET have very similar performance with the deterministic variant generating slightly better AUC values than the randomized variant, as expected. RESET provides measurably better classification accuracy than the other methods across all parameter settings except for the the low correlation case, for which VAM and GSVA have the best performance. Notably, ssGSEA generates nearly null AUC values of just slightly above 0.5 for this simulation design. As expected, performance of RESET improves with higher mean inflation, higher inter-gene correlation, larger gene set size and increased number of informative cells. Interestingly, VAM and GSVA performance decreases as inter-gene correlation is increased, which follows from both the impact of correlation on sparse count values (i.e., this will generate many cells with 0 values for most genes in the set) and the use by VAM of a correlation-breaking permutation to determine a null score distribution.

The results for the pure self-contained design, i.e., a model where all genes are included in the enriched set, are shown in Figure 2. It should be noted that the log-normalization process will largely eliminate the differential expression signature in this case, however, the higher initial mean for the impacted cells will lead to lower sparsity even after normalization and this is the pattern that is captured by VAM. For this simulation model, the relationship between RESET and VAM peformance is similar to that found for the block design with the gap between the methods just slightly larger, which makes sense given that VAM is a self-contained method so is insensitive to genes outside of the set. Of note, both GSVA and ssGSEA have nearly null performance for this model, which is expected given the competitive aspect of these techniques, i.e., they compare values for genes in the set to genes not in the set for a single cell. Although this type of data structure does not reflect a realistic biological scenario, it helps highlight the self-contained vs. competitive attributes of the various methods.

The results for the pure competitive design, i.e., a model where genes in the set are enriched for all cells, are shown in Figure 3. While the block and self-contained models could be assessed using only the scores for a single gene set, the pure competitive case requires comparison between the scores for the enriched set and the scores for non-enriched sets (four non-enriched sets were used for this simulation). The results for this model are quite dramatic, with RESET providing good classification accuracy across all parameter values, VAM yielding null values and both GSVA and ssGSEA generating AUC values significantly below 0.5. For VAM, the null performance is expected given the self-contained nature of the test, i.e., since all cells are enriched for the gene set, there is no self-contained signature to detect. For GSVA and ssGSEA, the very low AUC values are surprising. If a classifier consistently generated low AUC values, it would be feasible to turn it into a high AUC classifier simply by inverting the score ordering. In this case, however, GSVA and ssGSEA are generating both high and low AUC values for different scenarios so attempting to improve performance for the pure competitive design would break performance for the block design. In general, the scores generated by GSVA and ssGSEA on different sets are not directly comparable, i.e., they can be on very different scales for sets with largely the same pattern of expression. While standardization of the scores for each set can address this feature, such standardization would prevent detection of a pure competitive pattern.

### 3.2 Overall classification performance

We also evaluated RESET according to overall classification accuracy, i.e., can the method generate high overall scores for sets that have differential expression/correlation in a subset of the samples? The results for this evaluation are visualized in Figure 4. Because only RESET provides both overall and sample-level scores, just the RESET.det and RESET.ran variants are shown. Although it is challenging to intepret these results given the lack of comparative methods, they demonstrate very accurate performance for this simulation design across almost all parameter values.

**Figure 4:**
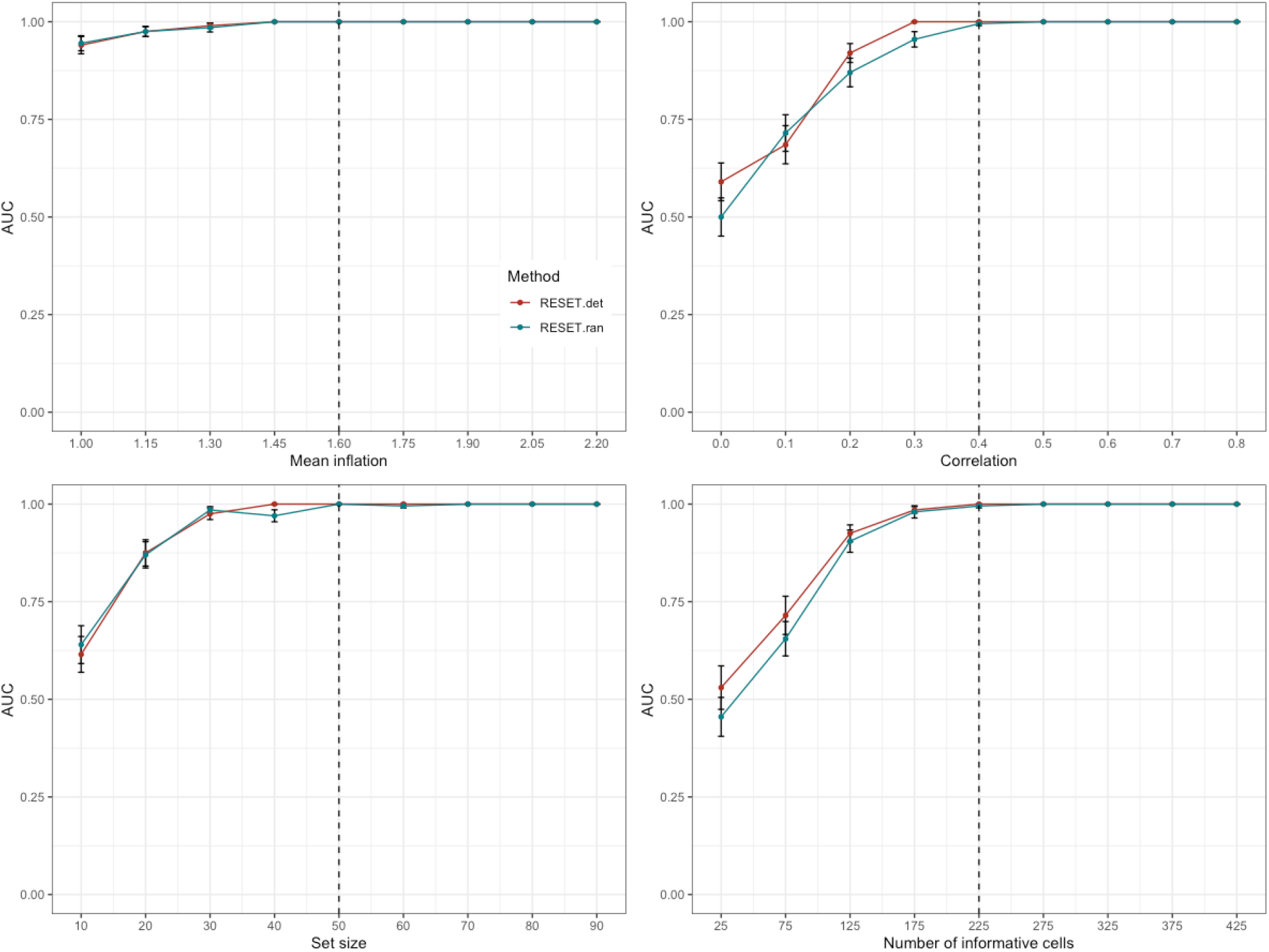
Overall classification performance of RESET.det and RESET.ran on scRNA-seq data simulated according to Section 2.7. Each panel illustrates the relationship between the area under the receiver operating characteristic curve (AUC) and one of the simulation parameters. The vertical dotted lines mark the default parameter value used in the other panels. Error bars represent the standard error of the mean.

### 3.3 Computational efficiency

Table 3.3 displays the relative execution time of GSVA, ssGSEA and VAM as compared to RESET. Relative times are shown for the analysis of the simulated data sets (2,000 cells and 500 genes) used to generate the classification results shown in Figure 1 (see Figure 5 below for performance on this simulated data as a function of gene set size), for the analysis of the 3k cell PBMC scRNA-seq data set using the the BioCarta (C2.CP.BIOCARTA) collection from the Molecular Signatures Database (MSigDB) [19] (see the *Human PBMC analysis* Section 3.4 for detailed results on the PBMC data set), for the analysis of the 11.8k cell mouse brain scRNA-seq data set using the MSigDB Gene Ontology biological process (C5.BP) pathway collection (see the *Mouse brain data analysis* Section 3.5 for detailed results on the mouse brain data set), and for the analysis of the very large 242k cell Mouse Cell Atlas (MCA) [37] scRNA-seq data set using a single gene set containing the first 50 genes. Since the R implementation of GSVA and ssGSEA force the conversion of the gene expression matrix into a non-sparse format, memory limitations prevented execution of these methods on the MCA data. For more details on the PBMC, mouse brain and MCA data sets and processing pipeline, see Section 2.8.

**Figure 5:**
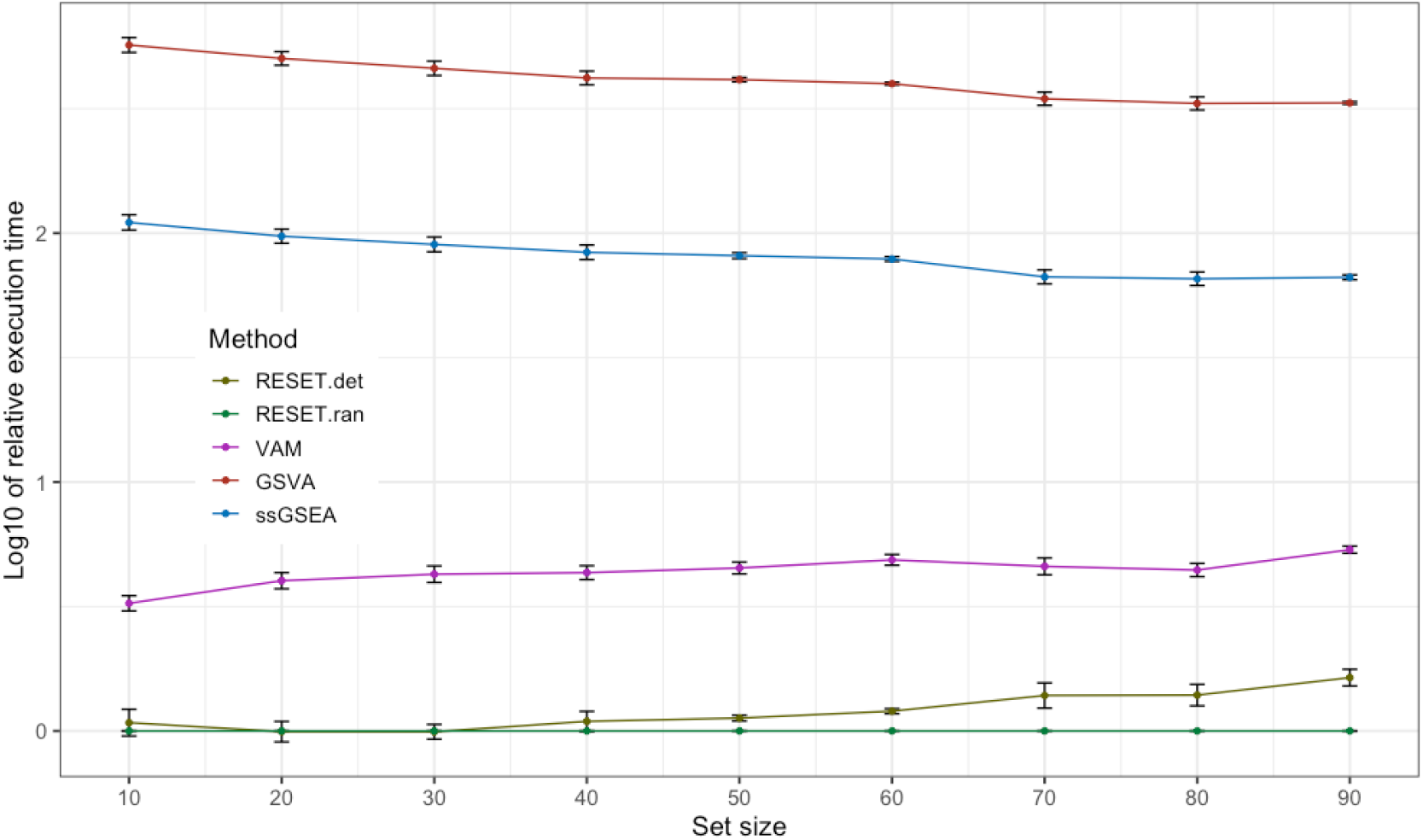
Average execution time of RESET.det, VAM, GSVA, and ssGSEA relative to RESET.ran. Relative values are plotted on the log_10_ scale. Execution times were computed on data simulated according to the procedure outlined in Section 2.7 for the block design. Error bars represent the standard error of the mean.

Across both the simulated and real scRNA-seq data, the RESET technique is two to four times as fast as VAM and nearly two to three orders-of-magnitude faster than GSVA and ssGSEA. As shown in Figure 5, the randomized version of RESET only provides a noticable performance benefit relative to the deterministic version of RESET when the gene set size is roughly four to five times larger than the target rank *k*.

### 3.4 Human PBMC analysis

As detailed in Section 2.8, we applied the RESET method and the comparison techniques to the 10x 2.7k human PBMC scRNA-seq data set. For this analysis, we looked at both the overall and cell-specific pathway scores generated by RESET. Table 2 lists the top 20 BioCarta pathways according to the overall RESET score, which accurately reflect the immune cell source of this scRNA-seq data set.

**Table 1:**
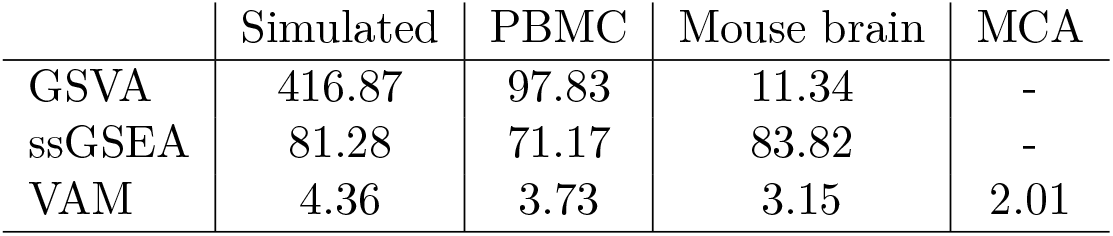
Relative execution time as compared to the RESET method on simulated scRNA-seq data, the PBMC scRNA-seq data set for MSigDB C2.CP.BIOCARTA collection, the mouse brain scRNA-seq data set for the MSigDB C5.BP collection, and the Mouse Cell Atlas for a single synthetic gene set. For the real scRNA-seq data, RESET was executed using the parameters specified in Section 2.8. For the simulated data, execution times are relative to the fully randomized version (i.e., ”RESET.ran”) as detailed in Section 2.7.

|  | Simulated | PBMC | Mouse brain | MCA |
| --- | --- | --- | --- | --- |
| GSVA | 416.87 | 97.83 | 11.34 | - |
| ssGSEA | 81.28 | 71.17 | 83.82 | - |
| VAM | 4.36 | 3.73 | 3.15 | 2.01 |

**Table 2:**
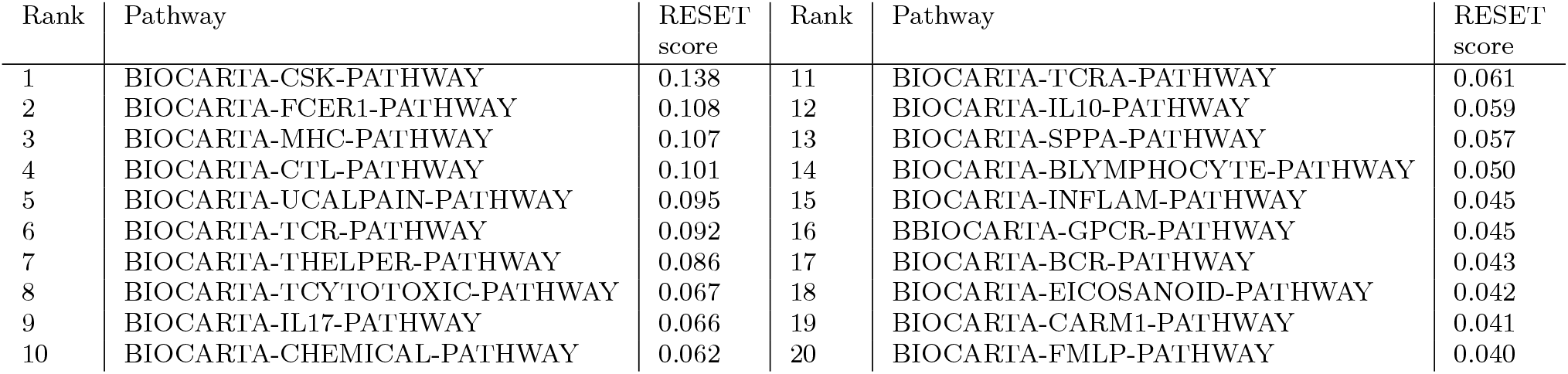
Top 20 BioCarta pathways according to overall RESET score for the PBMC data set

A important application of the cell-level scores computed by RESET involves the identification and visualization of differential pathway activity. Figure 6 illustrates such a visualization for the five BioCarta pathways most enriched in each cell type cluster according to the log2 fold-change in the mean RESET score of cells in the cluster relative to cells not in the cluster. These results provide important information regarding the range of pathway activity across all profiled cells. While many of the pathways shown in Figure 6 align the expected biology for the associated cell type, e.g., the B cell antigen receptor signaling pathway (as represented by the BIOCARTA-BCR-PATHWAY) has elevated scores in B cells, some of the results are unexpected, e.g, CD14+ monocytes have the highest scores for the BIOCARTA-TCR-PATHWAY. To correctly interprete the cell-level RESET scores, it is important to remember how the scores are computed mathematically and what that mathematical definition indicates about the structure of the analyzed scRNA-seq data. In particular, RESET scores capture how well a reduced rank representation of a given gene set can reconstruct a reduced rank representation of the entire data set. High cell-level RESET scores indicate that the value of set genes for a given cell can effectively reconstruct the values of all genes for that cell. As illustrated by the simulation studies, this can capture patterns of differential expression, however, it can also identify correlation patterns independent of any mean difference.

**Figure 6:**
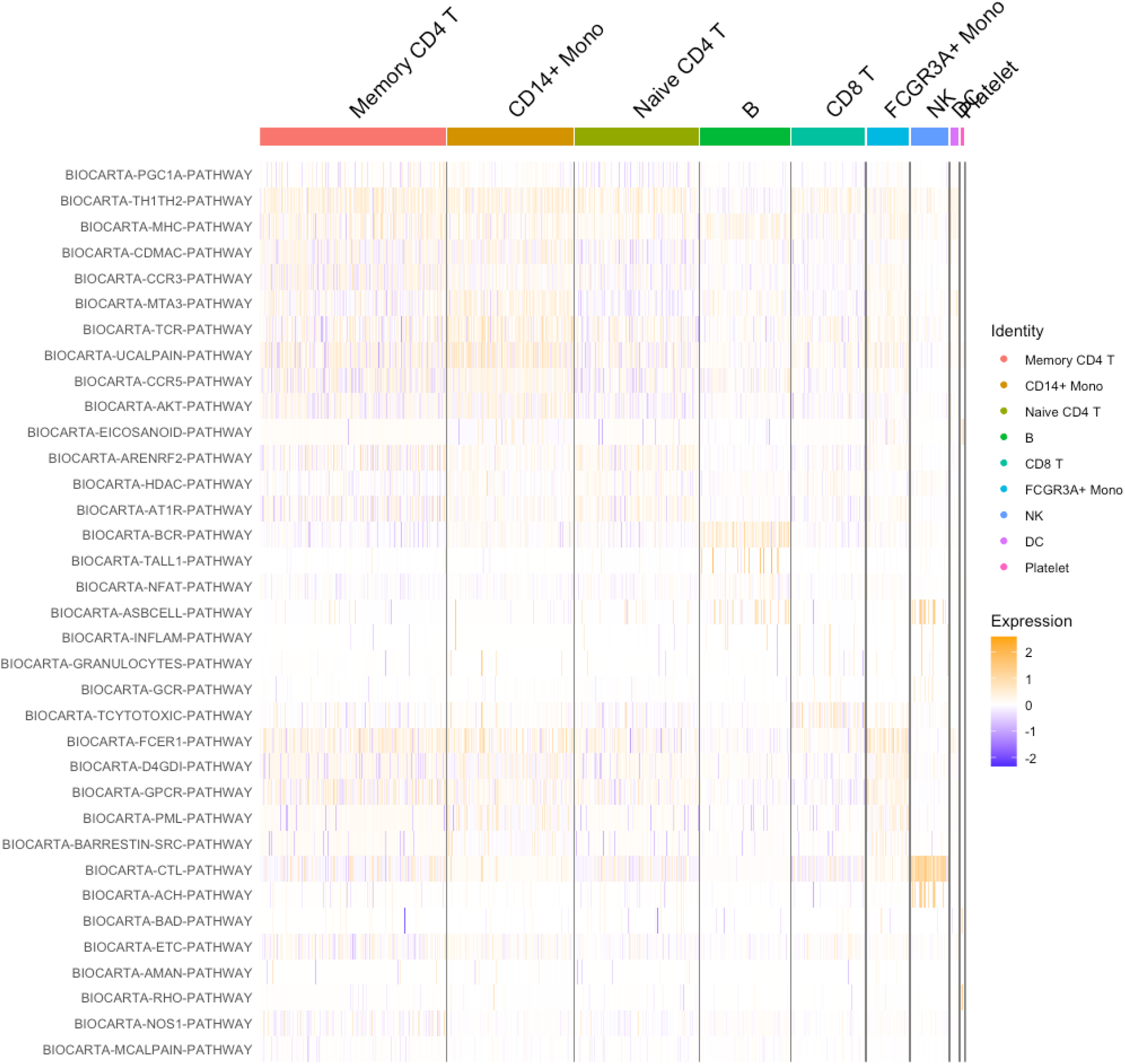
Heatmap visualization of the RESET cell-specific scores for the top five BioCarta pathways most enriched in each cluster of the PBMC scRNA-seq data according to the log2 fold-change in the mean RESET score of cells in the cluster relative to cells not in the cluster. Note that gene sets only appear once in the heatmap even if they are among the top five sets for multiple clusters.

Existing single sample gene set scoring methods like VAM, on the other hand, capture differences in mean expression. For such methods, high gene set scores will correspond to cells where mean expression of the genes in the set is elevated relative other cells in the data set. In general, RESET will produce distinct results from mean difference techniques and, for many use cases, performing both types of single sample gene set testing will provide the most comprehensive characterization of the data. Figure 7 illustrates the distinct results generated by RESET and VAM for four of the cells types in the PBMC data (note that the “BIOCARTA-” prefix has been removed to save space). While some pathways appear enriched using either RESET or VAM scores, e.g., BCR FOR B cells, TCYTOTOXIC for CD8 T cells, and CTL (Cytotoxic T Lymphocytes) for NK cells, others are enriched according to just one of the techniques. Biologically relevant pathways that are only enriched according to RESET scores include the ARENRF2 (Oxidative Stress Induced Gene Expression Via Nrf2) [38] and AT1R (Angiotensin II mediated activation of JNK Pathway via Pyk2 dependent signaling) [39] pathways for Naive CD4 T cells, the NFAT (NFAT and Hypertrophy of the heart) [40] and TALL1 (TACI and BCMA stimulation of B cell immune responses) pathways for B cells, the INFLAM (Cytokines and Inflammatory Response) and GRANULOCYTES (Adhesion and Diapedesis of Granulocytes) pathways for CD8 T cells, and the INFLAM and ASBCELL (Antigen Dependent B Cell Activation) [41] pathways for NK cells.

**Figure 7:**
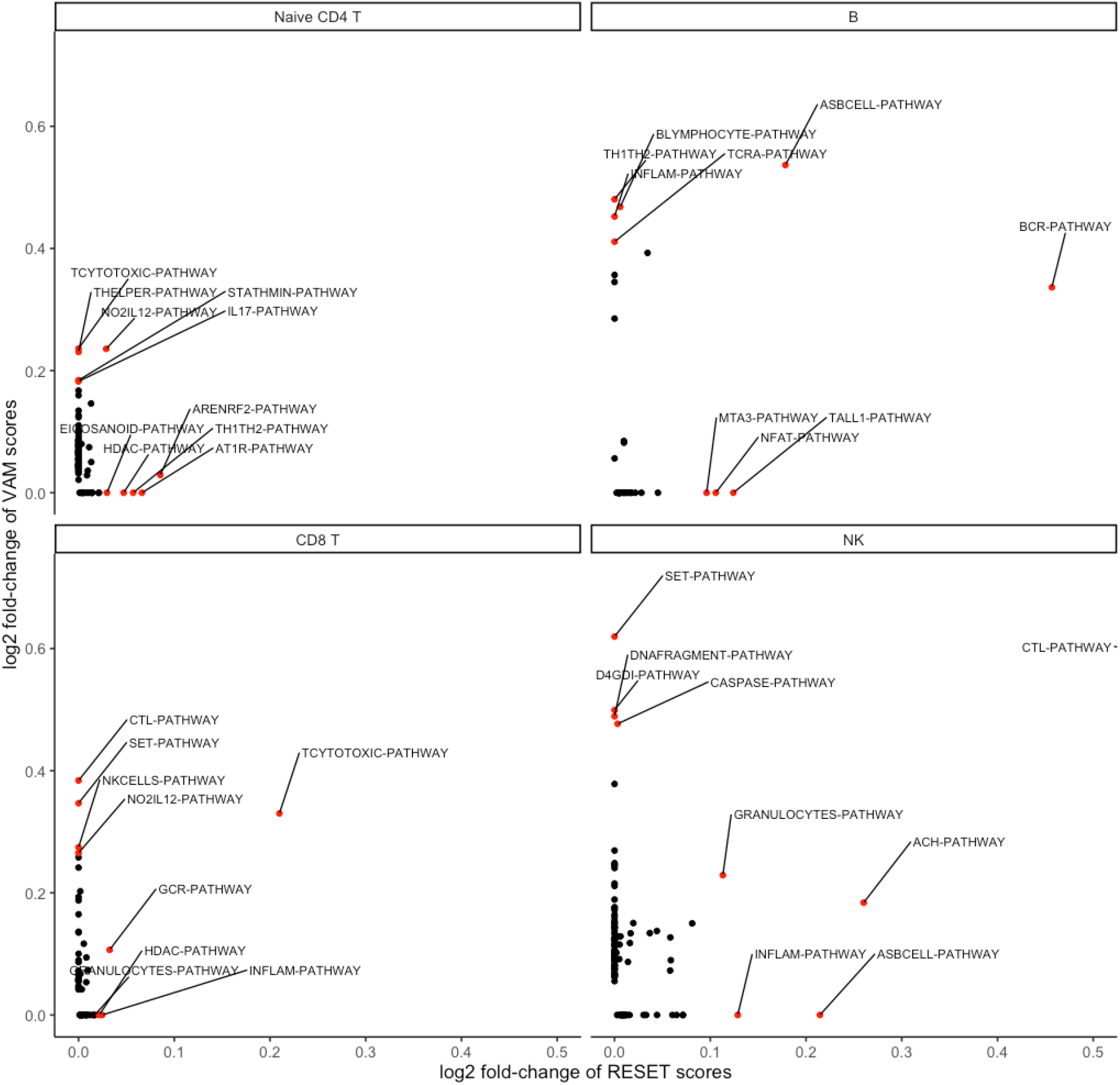
Visualization of cell type pathway enrichment as computed using either VAM or RESET scores.

### 3.5 Mouse brain cell analysis

As detailed in Section 2.8, we also analyzed the 10x 11.8k mouse brain scRNA-seq data set using RE-SET and the comparison techniques. For the mouse brain data, we used the SCTransform normalization technique instead of log-normalization and explored a much larger pathway collection (the MSigDB Gene Ontology (GO) biological process (C5.BP) collection). Figure 8 is a projection of the 9,320 cells remaining after quality control onto the first two UMAP dimensions with labels based on unsupervised clustering results and Figure 9 is a visualization of pathway enrichment in the six distinct clusters (2, 7, 9, 10, 11, and 12) according both RESET and VAM scores. In contrast to the PBMC analysis, RESET and VAM scores generated more similar pathway enrichment results for the mouse brain data. Importantly, prioritizing GO terms that appear enriched according to both methods can help identify the likely neuronal cell populations represented by the clusters: gabaeric interneurons for cluster 2, oligodentrocytes for cluster 7, granule cells for cluster 10, vascular cells for cluster 11 and microglial cells for cluster 12.

**Figure 8:**
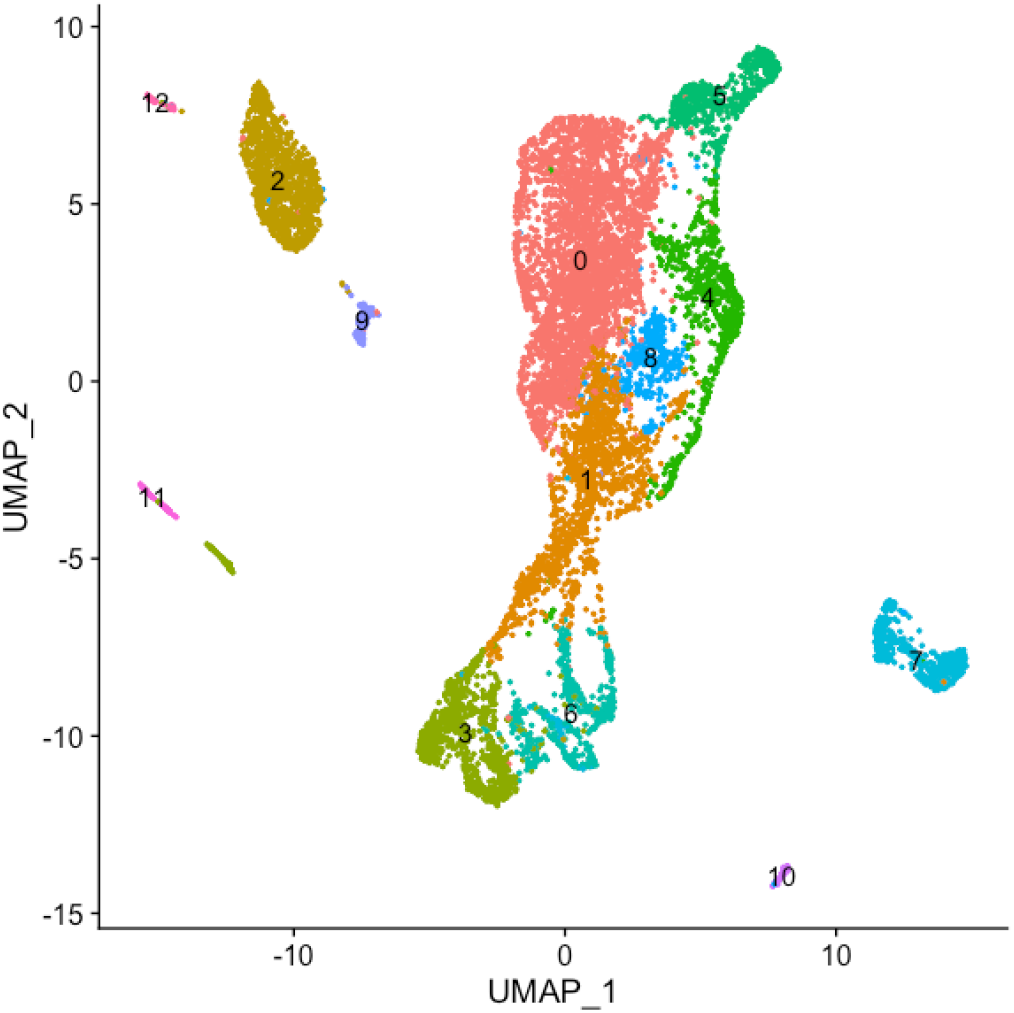
Projection of mouse brain scRNA-seq data onto the first two UMAP dimensions. Each point in the plot represents one cell, which are colored and labeled accounting to the output from unsupervised clustering.

**Figure 9:**
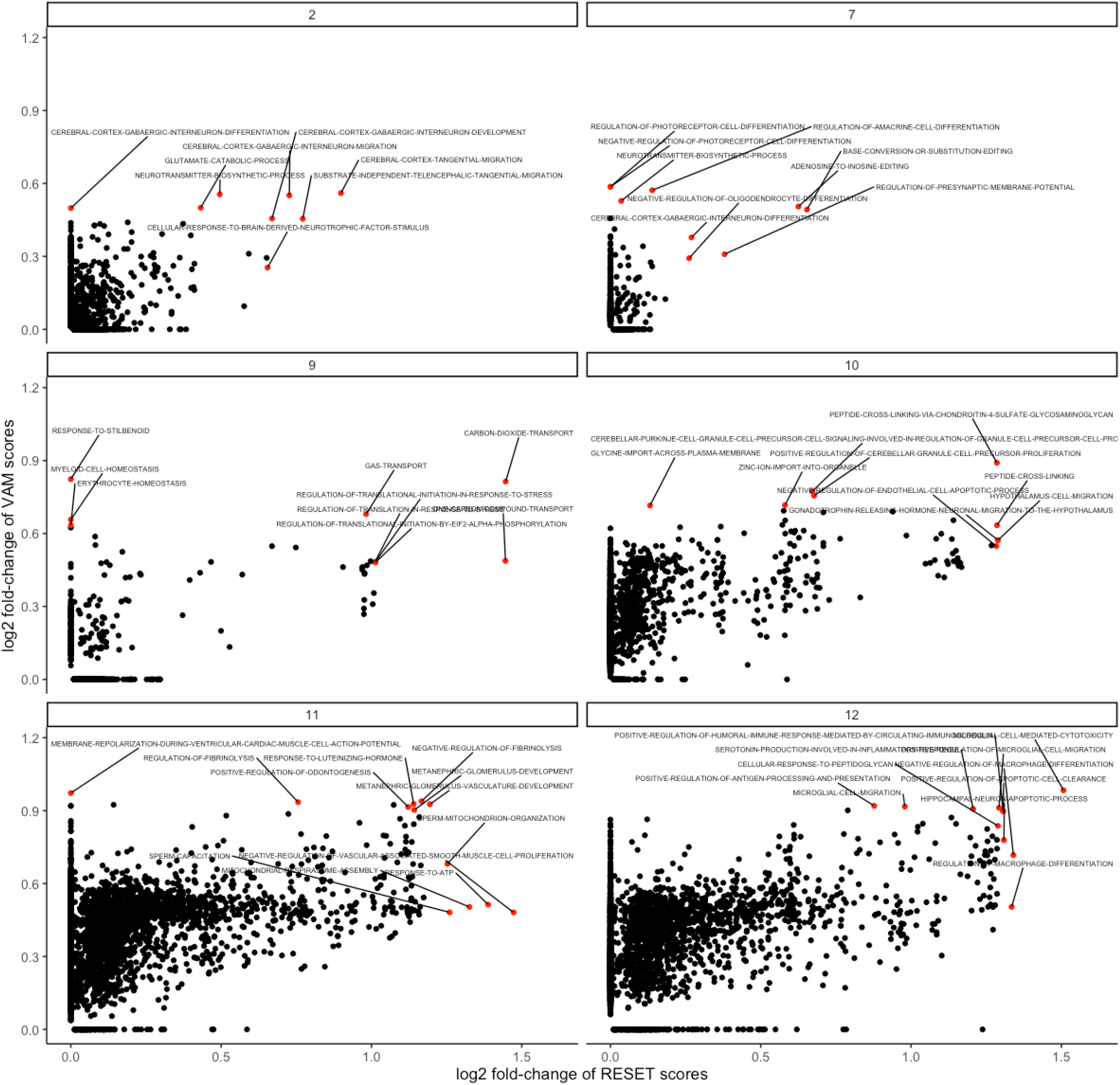
Visualization of cluster-level Gene Ontology biological process term enrichment as computed using either VAM or RESET scores.

